# Repairing neural damage in a *C. elegans* chemosensory circuit using genetically engineered synapses

**DOI:** 10.1101/2020.04.16.045443

**Authors:** Ithai Rabinowitch, Bishal Upadhyaya, Aaradhya Pant, Jihong Bai

## Abstract

Neuronal loss can considerably diminish neural circuit function, impairing normal behavior by disrupting information flow in the circuit. We reasoned that by rerouting the flow of information in the damaged circuit it may be possible to offset these negative outcomes. We examined this possibility using the well-characterized chemosensory circuit of the nematode worm *C. elegans*. In this circuit, a main sensory neuron class sends parallel outputs to several interneuron classes. We found that the removal of one of these interneuron classes impairs chemotaxis to attractive odors, revealing a prominent path for information flow in the circuit. To alleviate these deficiencies, we sought to reinforce a remaining neural pathway. We used genetically engineered electrical synapses for this purpose, and observed the successful recovery of chemotaxis performance. However, we were surprised to find that the recovery was largely mediated by inadvertently formed left-right lateral electrical connections within individual neuron classes. Our analysis suggests that these additional electrical synapses help restore circuit function by amplifying weakened neuronal signals in the damaged circuit. These results demonstrate the power of genetically engineered synapses to regulate information flow and signal intensity in damaged neural circuits.

## Introduction

The pattern of synaptic connectivity between neurons delineates possible paths for information flow in neural circuits. Loss of neurons due to injury or disease often causes a break in information flow, leading to behavioral deficiencies^1,2^. Remarkably, in some cases, the brain is capable of spontaneous recovery from such damage through mechanisms of synaptic plasticity^3,4^, suggesting that alternative circuit configurations could replace the original damaged ones and that information may be rerouted through alternative neural pathways, restoring signal propagation in the circuit. These observations raise questions as to how changes in synaptic connectivity impact information flow and whether such changes are sufficient for functional circuit recovery.

Here we address these questions by focusing on a well described neural circuit for chemosensation in the nematode worm *C. elegans*^5–7^. In this circuit, sensory information is transmitted from sensory neurons to a set of interneurons that modulate locomotion, enabling worms to locate chemical cues in the environment. We analyze the impact of neuronal loss on circuit function, and apply a synaptic engineering approach, based on the genetic insertion of new electrical synapses into the circuit^8,9^, to artificially reconfigure circuit connectivity, and thus, to construct a synaptic bypass designed to reinforce direct signal transmission from sensory neurons to undamaged interneurons. Our findings demonstrate the potential of synthetic electrical synapses to act as bypasses for neuronal information flow and as general amplifiers of weak signaling in damaged neural circuits.

## Results

### Loss of AIA interneurons impairs chemotaxis and AIB functionality

In the relatively simple and well-characterized *C. elegans* chemosensory neural circuit^5–7,10–13^, the AWC sensory neuron class sends parallel synaptic outputs to several interneuron classes, including AIA, AIB, and AIY (Fig. 1a). These interneurons act to modulate locomotion in an odor-dependent manner^5–7^. Together with additional premotor interneurons and motor neurons, the circuit enables worms to detect attractive odors and migrate towards them. Loss of AIA, one of the interneurons in the circuit, has been shown to adversely affect several behavioral capacities, such as salt chemotaxis learning^14^ and the integration of conflicting environmental cues^15^. In addition to these, we found, following genetic ablation of AIA, a reduction in chemotaxis to AWC-sensed odors such as isoamyl alcohol (IAA) and benzaldehyde (Fig. 1b). According to the *C. elegans* synaptic wiring diagram^16–18^ a major fraction of AIA synaptic outputs is directed to the AIB neuron class (Fig. 1a, inset). We thus wondered whether loss of AIA affects AIB function, and performed calcium imaging experiments to examine the responses of AWC, AIA, and AIB neurons to odor removal and presentation (Fig. 1c). As previously described^7^, odor removal in wild type worms triggered the activation of AWC, leading to AIA silencing and AIB activation (Fig. 1c, top). Conversely, odor presentation resulted in a drop in AWC activity, which produced a transient rise in AIA activity and a decrease in AIB activity (Fig. 1c, bottom). These response patterns have been attributed to inhibitory AWC→AIA and excitatory AWC→AIB synaptic transmission, respectively^7,19^ (Fig. 1a). Since AIB receives input from both AWC and AIA, we examined the extent of AIA impact on AIB activity. We found that AIA removal strongly diminishes AIB responses, suggesting that, in addition to direct AWC→AIB transmission, the indirect AWC→AIA→AIB pathway acts as a prominent driver of AIB activity (Fig. 1a). Removal of AIB in worms lacking AIA led to a further decrease in chemotaxis performance compared to AIA removal alone (Fig. 1d), suggesting that AIB still maintains residual functionality even in the absence of AIA, most likely due to the direct inputs from AWC.

**Figure 1.**
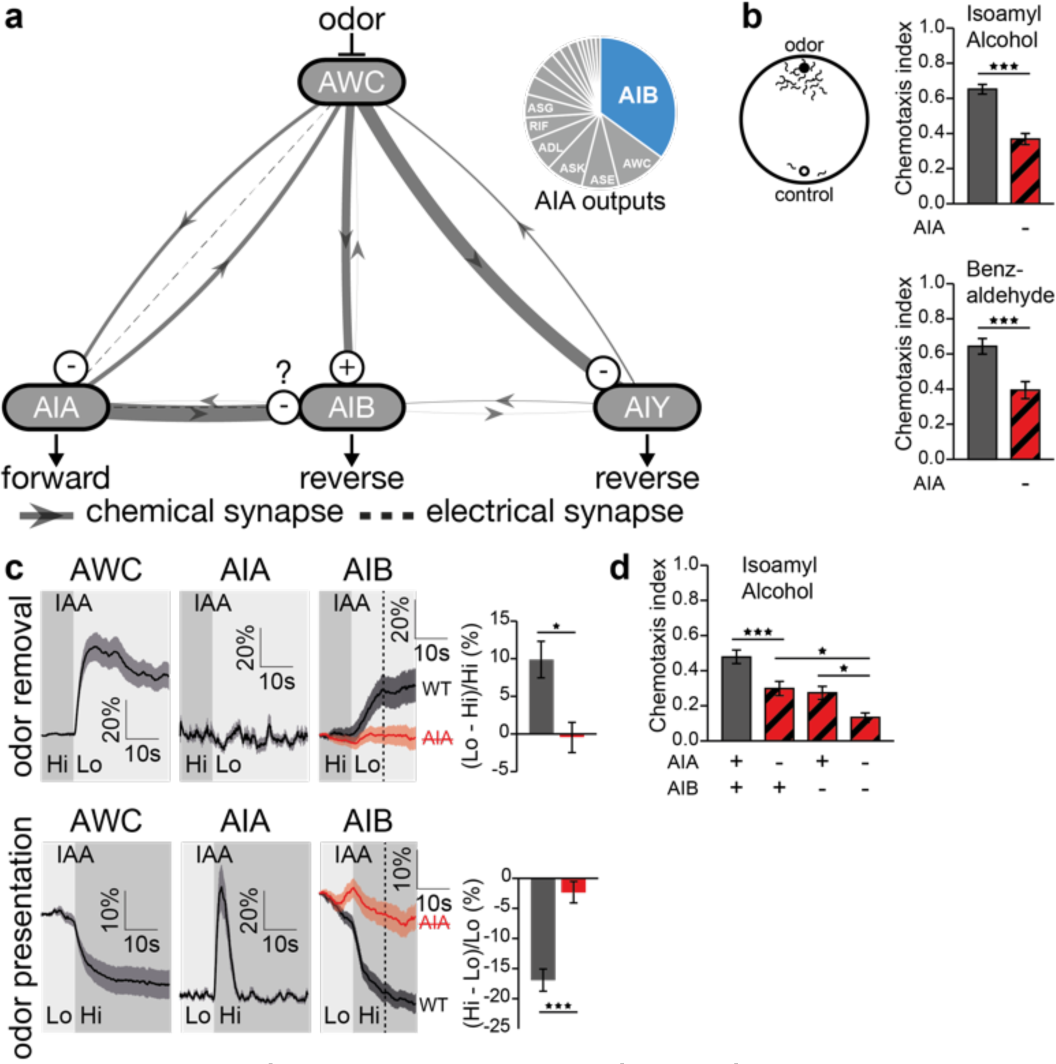
AIA plays an important role in chemosensory circuit function. (**a**) Simplified chemosensory circuit diagram. Edge thickness is proportional to connection strength (number of electron microscope sections that they occupy). Inset, relative strengths of AIA synaptic outputs. Source: www.wormwiring.org. (**b**) Impact of AIA removal on chemotaxis to AWC-sensed odors, isoamyl alcohol (IAA; n=40,40) and benzaldehyde (n=28,28). (**c**) Calcium responses of AWC, AIA and AIB neurons to removal and presentation of IAA. AIB responses (n=59,56 and n=90,87) are shown also for worms lacking the AIA neurons (red). (**d**) Impact of AIA and AIB removal on chemotaxis (n=65,63,64, 63). Error bars represent SEM. * p<0.05, ** p<0.01, *** p<0.001.

### Ectopic expression of connexin in AWC and AIB enhances chemotaxis in worms lacking AIA

We next asked whether reinforcing direct AWC→AIB transmission (Fig. 2a) could compensate for interrupted indirect AWC→AIA→AIB information flow in worms lacking AIA, thus enabling functional recovery of the circuit. To increase AWC and AIB coupling, we inserted a synthetic electrical synapse between the two neurons. We did this by ectopically expressing the vertebrate gap junction protein, connexin^20^, in each neuron (Fig. 2a). Connexin forms gap junctions that allow the passage of electrical signals between adjacent neurons (Fig. 2a, inset). Vertebrate connexin ectopically expressed in *C. elegans* neurons, is unlikely to interact with the endogenous invertebrate gap junction protein, innexin^21^, and is thus expected to give rise to highly specific new connections. We have previously established and validated this technique under different circuits and scenarios^8,22^. Upon testing worms that express connexin in both AWC and AIB, we observed a substantive recovery of chemotaxis performance, exceededing even that of wild type worms (Fig. 2b). We also expressed connexin in AWC and AIB separately as controls, and were surprised to find that each of these was sufficient for restorating chemotaxis, equivalent in degree to that of the normal worms (Fig. 2b).

**Figure 2.**
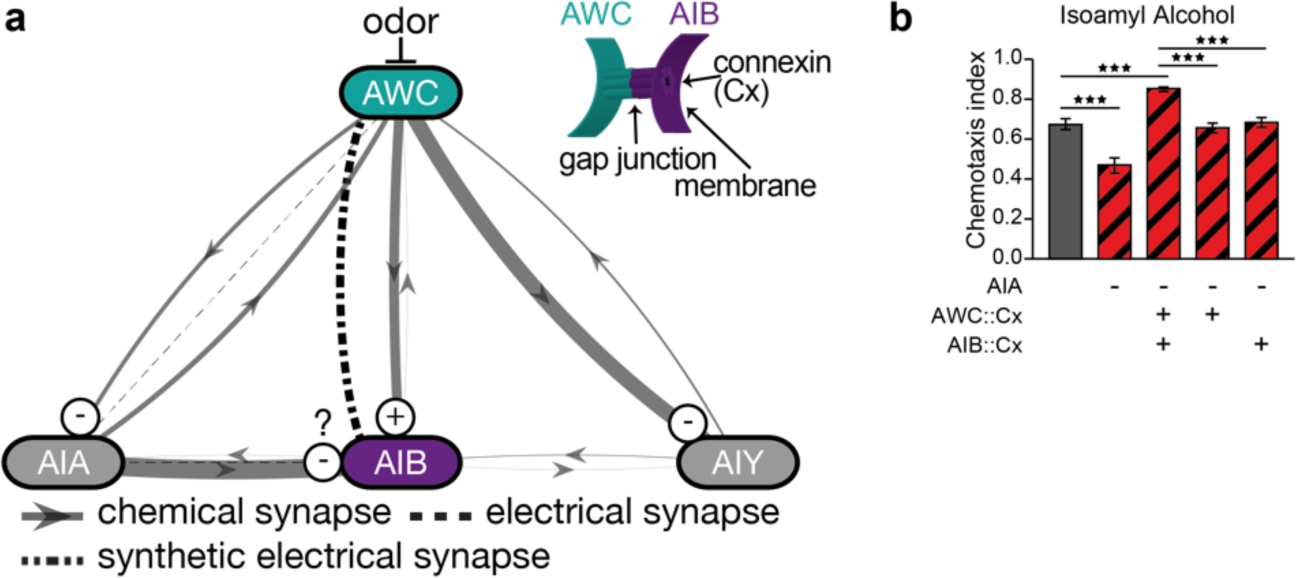
Ectopic expression of connexin in AWC and AIB. (**a**) Circuit diagram showing a desired new link between AWC and AIB formed by an engineered electrical synapse. Inset, the synthetic AWC-AIB electrical synapse, or gap junction, is constructed by expressing connexin in the AWC and AIB. (**b**) Chemotaxis following AIA removal and ectopic expression of connexin in AWC alone, AIB alone or both (all n=21). Error bars represent SEM. *** p<0.001.

### Connexin forms synthetic electrical synapses between left-right neuron pairs within individual neuron classes

How could connexin expression in a single neuron class induce functional recovery in the damaged circuit? To address this question, we considered that many *C. elegans* neuron classes, including those in the olfactory circuit, are composed of bilateral left-right neuron pairs^23^ (Fig. 3a). Connexin expression in these neurons could thus lead to the formation of new electrical synapses between the left and right members of the class (Fig. 3a). A sensible test of this possibility is to activate just one of the two individual neurons within a neuron pair and then examine whether connexin expression in both neurons equalizes their responses due to synthetic electrical coupling. To test this, we examined the AIB neuron pair. No synaptic connections are known to exist between AIBL and AIBR (Fig. 3a), although their two processes pass in close proximity to each other in their paths across the nerve ring^16^. We sought to test whether connexin expression in the AIB pair results in the formation of synthetic electrical coupling between these neurons. We noticed an asymmetry in the connection strength between the salt sensing neuron, ASER, and the two AIB neurons, AIBL and AIBR (Fig. 3a). We asked whether this asymmetry in connectivity could lead to distinct AIBL and AIBR responses to ASER input. To examine this possibility, we activated ASER by exposing worms to a sudden decrease in salt concentration^11^, and recorded AIBL and AIBR responses. Such ASER activation has been shown to elicit a rise in the average AIB calcium response^24^. Notabley, when we analyzed the left and right AIB neuron responses separately, we were able to observe an asymmetrical response to this stimulation: whereas AIBL strongly increased its activity, AIBR exhibited almost no response (Fig. 3b). We took advantage of this lateralization that we observed in the AIB salt response, and tested whether connexin expression in AIB would diminish this asymmetry. Indeed, this led to a much more similar response in AIBL and AIBR to the decrease in salt concentration. This result supports the idea that connexin expression in AIB, creates functional electrical coupling between AIBL and AIBR.

**Figure 3.**
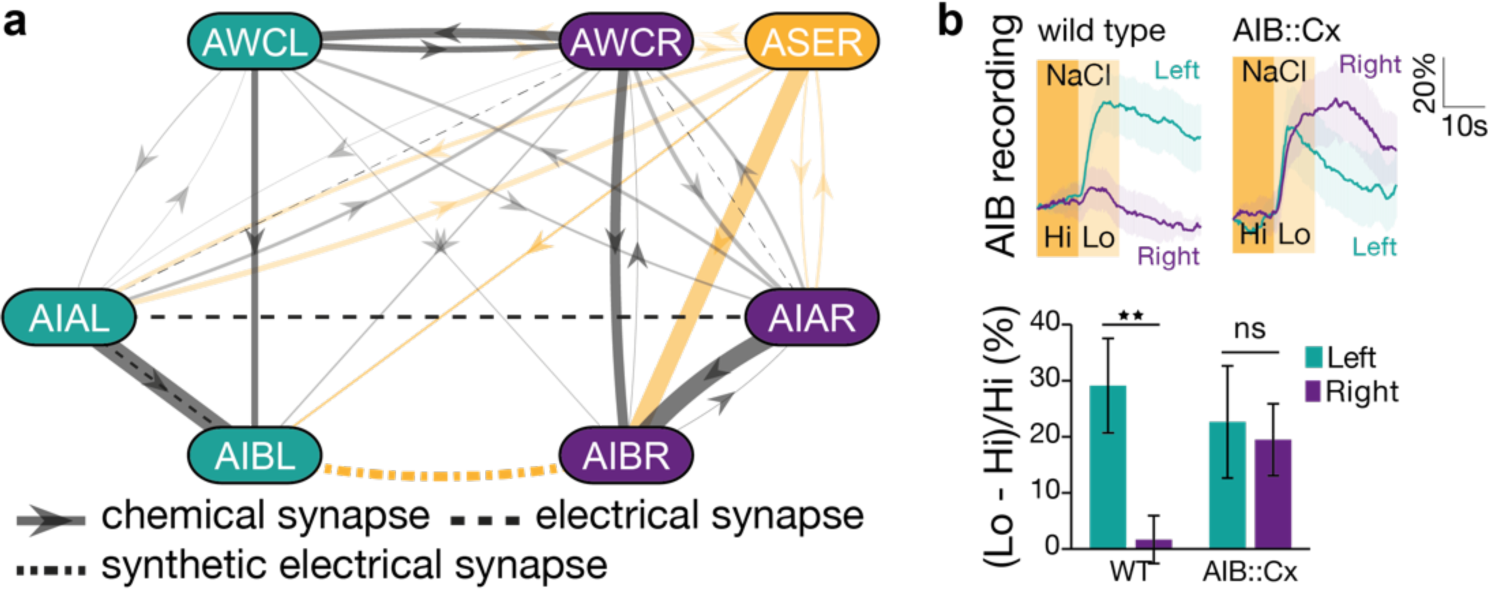
Synthetic electrical coupling between left-right neuron pairs. (**a**) Circuit diagram showing individual neurons in the olfactory circuit from each bilateral symmetrical neuron class. In addition, an asymmetrical distribution of synaptic strengths between ASER and AIBR versus AIBL is shown. (**b**) Activation of ASER by reducing salt concentration, led to strong activation of AIBL and almost no response in AIBR. Ectopic connexin expression in AIB, equalized the response of the two neurons, as expected for an AIBL-AIBR electrical coupling (n=28,25,25,25). Error bars represent SEM. ** p<0.01.

### Lateral electrical coupling can amplify weak signals

How could artificial left-right electrical coupling lead to restored chemotaxis in worms lacking AIA? Previous studies in the olfactory systems of drosophila^25,26^ and zebrafish^27^ have identified a role for electrical synapses in amplifying weak signals in these circuits through lateral excitation^28^. We asked whether the synthetic left-right electrical coupling obtained through connexin expression within individual neuron classes could artificially amplify weak signals in the *C. elegans* olfactory circuit. To test this, we first studied the responses of AWC neurons to weak stimulation using low IAA concentrations. As expected, reduced odor intensity led to a decrease in the magnitude of AWC responses in normal worms (Fig. 4a, cf. 1:10^−5^ and 1:10^−7^). This weakened response was considerably amplified in worms expressing connexin in AWC (Fig. 4a, top). Connexin expression in AWC produced also a corresponding behavioral effect, substantially increasing chemotaxis to dilute IAA, compared to normal worms (Fig. 4a, bottom). These findings suggest that electrical coupling between left and right AWC neurons can amplify weak signals.

**Figure 4.**
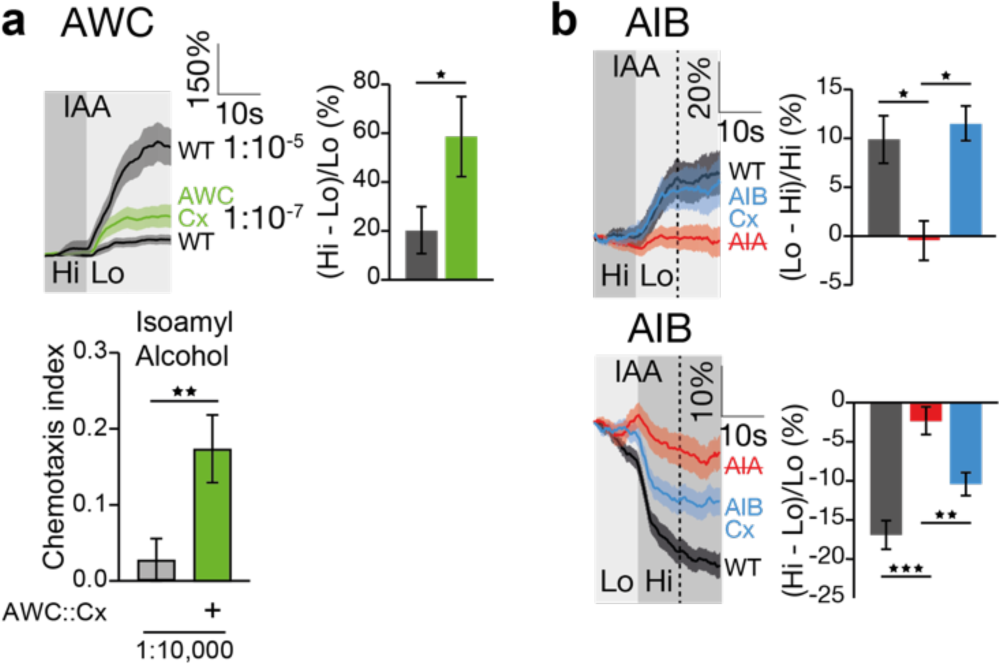
Left-right electrical coupling can amplify weak signals. (**a**) AWC responses to IAA removal considerably weaken when IAA concentration is decreased. However, ectopic expression of connexin in AWC increases the weakened AWC response (top; n=24,24) and enhanced chemotaxis to dilute IAA (bottom; n=26,23). (**b**) AIB weakened responses following AIA removal (n=59,56,61) and presentation (n=90,87,95) are partially restored by ectopic expression of connexin in AIB. Error bars represent SEM. * p<0.05, ** p<0.01, *** p<0.001.

We next considered the impact of synthetic AIBL-AIBR coupling on signal strength. As we have shown above, AIA removal strongly decreases AIB calcium responses (Fig. 4b). We asked whether expressing connexin in AIB mitigated the weakened response, amplifying AIB signaling. We found that AIB responses to both AIA removal and presentation were almost completely restored in worms expressing connexin in their AIB neurons (Fig. 4b), which correlated with the recovery of chemotaxis behavior (Fig. 2b). Taken together, our results suggest that inserting synthetic electrical synapses between left and right neuron class members could amplify weak signals in these neurons and lead to a circuit-level recovery from damage.

### Reinforcing the direct AWC→AIB pathway contributes to functional recovery

Beyond the observed restorative effects of inserting synthetic electrical connections between left and right neuron pairs in the olfactory circuit, we sought to evaluate the impact of AWC-AIB electrical coupling on the direct AWC→AIB pathway (Fig. 2a). In particular, we wished to determine whether the high chemotaxis performance observed in worms expressing connexin in both AWC and AIB (Fig. 2b) was due, at least in part, to enhanced AWC-AIB coupling, or was solely the sum of the effects of independent connexin expression in AWC and AIB. To address this question, we engineered a synthetic electrical connection between just one of the two AWC neurons and AIB. We used the Pstr-2 promoter, specific to AWC^ON^ (as well as the ASI neurons^29^), to selectively drive connexin expression in AWC^ON^ but not AWC^OFF^, in worms lacking AIA. These worms are not expected to display chemotaxis recovery because they do not form AWCL-AWCR connections. Indeed, no such recovery was observed (Fig. 5a). To the contrary, the defects in chemotaxis were even further exacerbated (Fig. 5a). This could be due to the formation of additional new electrical synapses with or within another neuron class. We thus proceeded to examine the effects of combined expression of connexin in AWC^ON^ and AIB, and found a significant increase in chemotaxis performance beyond the levels obtained by connexin expression in AIB alone (Fig. 5b). These results are consistent with AWC^ON^-AIB synthetic coupling having a distinct role in functional recovery. It is worth noting that the observed effect may underestimate the full capacity of AWC-AIB synthetic coupling to bypass the break in information flow caused by the loss of AIA. First, the AWC^ON^-AIB connection includes only one of the two AWC neurons (AWC^ON^). It is thus possible that linking both AWC neurons to AIB could deliver enhanced stimulation to AIB with even greater impact. Second, as described (Fig. 5a), the AWC^ON^ connexin worms show diminished chemotaxis, which could counteract the positive contribution of AWC^ON^-AIB.

**Figure 5.**
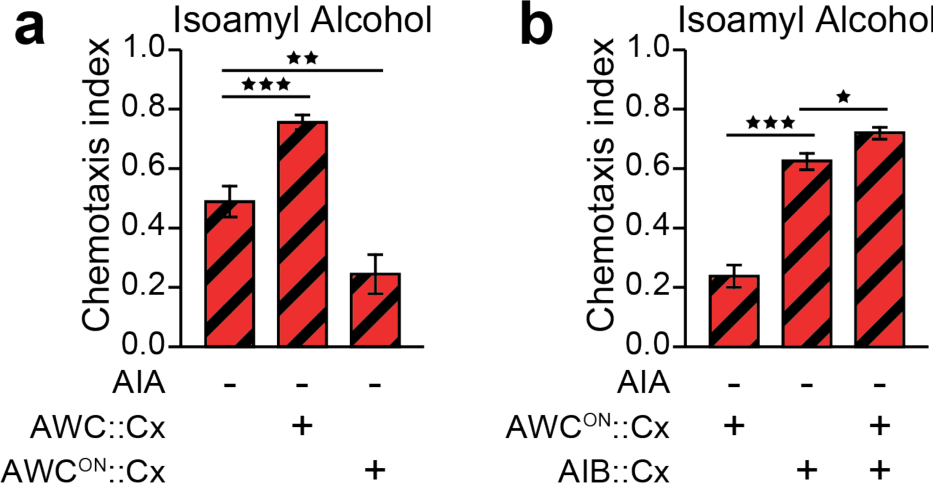
Direct synthetic coupling of AWC and AIB contributes to functional recovery in worms lacking AIA. (**a**) Ectopic connexin expression using an AWC^OFF^ promoter does not restore chemotaxis behavior (all n=18). (**b**) Ectopic expression of connexin in AWC^OFF^ and AIB results in an increased recovery compared to connexin expression in AIB only (all n=64). Error bars represent SEM. * p<0.05, ** p<0.01, *** p<0.001.

## Discussion

It is often convenient to describe information flow in the nervous system as originating from the sensory neurons, passing through interneurons, and terminating in motor neurons. The *C. elegans* olfactory circuit is frequently depicted in this manner (e.g. Fig. 1a): chemosensory information is transmitted from the AWC sensory neurons to several interneurons, including AIA and AIB, and then from there downstream to premotor and motor neurons. However, our findings support a more distributed rather than linear view of information flow in neural circuits, whereby an indirect pathway, such as AWC→AIA→AIB, appears to be more prominent than the direct AWC→AIB stream, and lateral electrical coupling within neuron classes such as AIBL-AIBR can regulate signal strength in the circuit. Indeed, it is tempting to speculate that one of the reasons why AIA plays such an essential role in the olfactory circuit is because of the electrical synapses that naturally couple AIAL and AIAR (Fig. 3a), and possibly provide this neuron class with a level of gain control.

The general architecture of the *C. elegans* olfactory circuit has previously been compared to that of retinal circuits of photoreceptor cells and their postsynaptic partners, the ON and OFF bipolar cells^7^. Our analysis reveals additional analogies with the retinal circuit. Similarly to the indirect functional links between AWC and AIB in the *C. elegans* olfactory circuit, also photoreceptor cells form indirect links with ON and OFF bipolar cells^30^. Furthermore, many neuron classes in the retina are interconnected by electrical synapses^31^. Such electrical coupling could play a role in signal amplification as in the olfactory circuit. This conserved feature of electrical synapses, shared with other organisms^25–27^, may reveal a potential general strategy for using synthetic electrical coupling as a means for signal amplification in damaged circuits.

The notion of creating artificial links to overcome neural circuit damage is proving to be especially effective in brain-machine-brain interfaces (BMBIs)^32^. These electronic devices composed of electrodes and computers, monitor and decode neural signals from one brain region, and then encode and deliver corresponding stimulation to a second region^33^. Such systems appear to be capable of bridging disconnected neural pathways, and thus restoring function after neural damage^34,35^. Our current work follows a similar logic, but offers a different, complementary implementation of this basic approach. It focuses on synthetic neuronal coupling at the microscopic level of small local circuits rather than general macroscopic brain regions linked by BMBIs. Thus, the design and implementation of genetically encoded synapses, could complement BMBIs in promoting novel strategies for coping with neuronal damage and for uncovering circuit pricinples.

## Methods

### *C. elegans* Growth and Maintenance

*C. elegans* strains were maintained under standard conditions at room temperature (20-21°C) on nematode growth medium (NGM) 2% agar plates seeded with *E. coli* strain, OP50. The N2 strain (Bristol, England) was used as the wild type reference.

### Genetic Engineering of Electrical Synapse

New electrical synaptic connections were inserted into the chemotaxis circuit by ectopically expressing the gene for connexin 36 (Cx36) in specific neurons, as previously described^8^. Briefly, a codon-optimized synthesized cDNA sequence of *Mus musculus* gap junction protein, delta 2 (Gjd2), known also as connexin 36 (Cx36), was fused to relevant upstream promoters (Podr-1 for AWC, Pinx-1 for AIB, Pstr-2 for AWC^ON^) and a downstream gene encoding mCherry, and inserted into a MosSCI vector (see below) using Gateway.

### Transgenic Strain Generation

Strains expressing genetically encoded calcium indicators were generated by standard microinjection of DNA constructs at 50-70 ng/µl together with specific co-injection markers at 10-30 ng/µl. Strains expressing connexin were generated using the MosSCI system for single copy targeted chromosomal integration^36^. The injection mix included pCFJ601 *Peft-3::mos1 transposase::tbb-2utr* at 50 ng/µl for cutting out the Mos1 transposon, a DNA repair template including the connexin construct at 30 ng/µl, and pMA122 *Phsp-16.41::peel-1::tbb-2 UTR* at 10 ng/µl for eliminating extrachromosomal transformants. To combine different transgenic strains we performed standard crossings. For unknown reasons, the cross between *peIs580* (genetically ablated AIA) and *pekSi47* (connexin expressed in AWC^ON^) yielded homozygotes only for *peIs580*, and so that the resultant crossed strain (BJH2170) that we used was heterozygotic for *pekSi47*.

| Strain | Purpose | Figure |
| --- | --- | --- |
| **JN580\*** *pels580[Pins-1(short)::casp1; Pins-1(short)::venus Punc-122::gfp]* | AIA genetic ablation | 1b,d; 2b; 5a,b |
| **AQ2632** *ljEx351[Podr-1::YC3.60\*, Pelt-2::mCherry]* | AWC calcium imaging | 1c |
| **CX11073\*** *kyEx2916[Pgcy-28.d::GCamp2.2b, Punc-122::gfp]* | AIA calcium imaging | 1c |
| **BJH1033** *pekEx176[Pinx-1::GCaMP6s, Punc122::mCherry, Pgcy-5::ChrimsonCel::SL2::mCherry]* | AIB calcium imaging | 1c; 3b; 4b |
| **BJH2190** *pels580[Pins-1(short)::casp1; Pins-1(short)::venus Punc-122::gfp]; pekEx176[Pinx-1::GCaMP6s, Punc122::mCherry, Pgcy-5::ChrimsonCel::SL2::mCherry]* | AIB calcium imaging, AIA ablated | 1c |
| **BJH2176** *pels580[Pins-1(short)::casp1; Pins-1(short)::venus Punc-122::gfp]; pekSi46[cb-unc-119(+), Podr-1::Cx36::mCherry, Pvha-6::mCherry] ttTi5605 II; pekSi54[cb-unc-119(+), Pinx-1::Cx36::mCherry, Pvha-6::GFP] cxTi10816 IV* | AIA ablated and connexin in AWC and AIB | 2b |
| **BJH2173** *pels580[Pins-1(short)::casp1; Pins-1(short)::venus Punc-122::gfp]; pekSi46[cb-unc-119(+), Podr-1::Cx36::mCherry, Pvha-6::mCherry] ttTi5605 II* | AIA ablated and connexin in AWC | 2b; 5a |
| **BJH2168** *pels580[Pins-1(short)::casp1; Pins-1(short)::venus Punc-122::gfp]; pekSi54[cb-unc-119(+), Pinx-1::Cx36::mCherry, Pvha-6::GFP] cxTi10816 IV* | AIA ablated and connexin in AIB | 2b; 5b |
| **BJH1055** *pekEx196[Podr-1::GCaMP6s-SL2-mCherry + Punc122::mCherry]* | AWC calcium imaging | 4a |
| **BJH2463** *pekEx196[Podr-1::GCaMP6s-SL2-mCherry + Punc122::mCherry]; pekSi46[cb-unc-119(+), Podr-1::Cx36::mCherry, Pvha-6::mCherry] ttTi5605 II* | AWC calcium imaging and connexin in AWC | 4a |
| **BJH2192** *pekSi54[cb-unc-119(+), Pinx-1::Cx36::mCherry, Pvha-6::GFP] cxTi10816 IV; pekEx176[Pinx-1::GCaMP6s, Punc122::mCherry, Pgcy-5::ChrimsonCel::SL2::mCherry]* | AIB calcium imaging and connexin in AIB | 3b; 4b |
| **BJH918** *pekSi46[cb-unc-119(+), Podr-1::Cx36::mCherry, Pvha-6::mCherry] ttTi5605 II* | Connexin in AWC | 4a |
| **BJH2170\*\*** *pels580[Pins-1(short)::casp1; Pins-1(short)::venus Punc-122::gfp]; pekSi47[cb-unc-119(+), Pstr-2::Cx36::mCherry, Pvha-6::mCherry] cxTi10816 IV* | AIA ablated and connexin in AWC<sup>ON</sup> | 5a,b |

★ Strain JN580 kind gift of Yuichi Iino. Strain CX11073 kind gift of Cori Bargmann.
★★ Strain BJH2170 is heterozygotic for *pekSi47*.

### Chemotaxis Assay

To evaluate chemotaxis, we performed standard behavioral assays^37^. Approximately 200 adult worms were washed with M9 buffer onto 9 cm NGM plates. A 1 µl drop of 1M sodium azide was placed at two spots on either side of the plate. A 10 µl drop of Isoamyl alcohol (IAA) or benzaldehyde (Bz) diluted in ethanol was applied at one spot and a 10 µl drop of ethanol was placed at the other spot as a control. We found that this relatively large volume of drops helped reduce variance. The plate was divided into three regions. The number of worms immobilized within the two regions containing the spots was counted and used to calculate the chemotaxis index (the difference in worm number between stimulus region and control region, divided by the sum of worms found in these two regions). Each experiment was repeated in at least 3 different days. The total number of repetitions for each condition is indicated in the figure legends. One-way ANOVA was performed to compare between multiple group means. T-tests were carried out for comparison between two specific groups. Bonferroni corrections were applied for multiple comparisons.

### Calcium Imaging

Calcium imaging was performed using a microfluidic PDMS chip designed to deliver chemical stimuli under a fluorescent microscope^38^. We placed the chip on top of a 63x oil objective of a Leica DMI3000B inverted microscope and a QImaging OptiMOS camera. 10 sec after the beginning of each recording, we switched on the stimulus channel presenting the worm with either odor (isoamyl alcohol, AIA) or salt, diluted in S-basal medium lacking cholesterol. Imaging continued for an additional 30 sec. We then kept the stimulus channel open for 5 min and resumed imaging. After an additional 10 sec, we switched the stimulus off and continued the recording for 30 sec. Images were captured at a rate of 5 frames per second with a 100ms exposure time. For each recording, a region of interest was defined as a square-shaped area surrounding the desired cell body. Background-subtracted fluorescence intensity values were collected from every sample’s ROI and stored into MATLAB formatted files. Changes in fluorescence intensity (ΔF/F%) were calculated by dividing each value by the average intensity of the first 3 seconds of imaging. For statistical analysis we calculated the ratio of the average fluorescence intensity over the ten seconds after and before stimulus presentation/removal. One-way ANOVA was performed to compare between multiple group means. T-tests were carried out for comparison between two specific groups. Bonferroni corrections were applied for multiple comparisons.

## Acknowledgments

This research was supported by Hartwell Innovation Fund, R21DC016158, and R01GM127857 to JB. The authors thank the Caenorhabditis Elegans Genetic Consortium (funded by NIH Office of Research Infrastructure Programs P40 OD010440) and Dr. Shohei Mitani, Dr. Yuichi Iino and Dr. Cornelia Bargmann for worm strains. We thank the following people for technical assistance (Lin Zhang, Norman Nguyen, Carli Wightman).

